# Predicting educational achievement from genomic measures and socioeconomic status

**DOI:** 10.1101/538108

**Authors:** Sophie von Stumm, Emily Smith-Woolley, Ziada Ayorech, Andrew McMillan, Kaili Rimfeld, Philip S. Dale, Robert Plomin

**Author notes:** **CORRESPONDING AUTHOR**, Sophie von Stumm at. **AUTHOR CONTRIBUTIONS**, S.v.S and R.P. conceived and designed the study. S.v.S. and A.M. analysed the data. S.v.S. and R.P. drafted the paper and E.S.-W., Z.A., A.M., K.R. and P.S.D. helped to revise the paper. All authors approved the final draft of the paper. **OPEN PRACTICES STATEMENT**, The analyses reported in this manuscript were not pre-registered. The data and materials are available from the permanent TEDS archive upon request to R.P.

## Abstract

The two best predictors of children’s educational achievement available from birth are parents’ socioeconomic status (SES) and, recently, children’s inherited DNA differences that can be aggregated in genome-wide polygenic scores (GPS). Here we chart for the first time the developmental interplay between these two predictors of educational achievement at ages 7, 11, 14 and 16 in a sample of almost 5,000 UK school children. We show that the prediction of educational achievement from both GPS and SES increases steadily throughout the school years. Using latent growth curve models, we find that GPS and SES not only predict educational achievement in the first grade but they also account for systematic changes in achievement across the school years. At the end of compulsory education at age 16, GPS and SES respectively predict 14% and 23% of the variance of educational achievement. Analyses of the extremes of GPS and SES highlight their influence and interplay: In children who have high GPS and come from high SES families, 77% go to university, whereas 21% of children with low GPS and from low SES backgrounds attend university. We find that the associations of GPS and SES with educational achievement are primarily additive, suggesting that their joint influence is particularly dramatic for children at the extreme ends of the distribution.

**HIGHLIGHTS:**

- Genome-wide polygenic scores (GPS) and socioeconomic status (SES) account together for 27% of the variance in educational achievement from age 7 through 16 years
- The predictive validity of GPS and SES increases over the course of compulsory schooling
- The association of GPS and SES is primarily additive: their joint long-term influence is particularly pronounced in children at the extreme ends of the distribution
- 77% of children with high GPS from high SES families go to university compared to 21% of children with low GPS from low SES

## INTRODUCTION

Until recently, measures of parents’ socioeconomic status (SES) have been the most powerful predictors available from birth of differences in the normal range of children’s educational achievement. Although typically operationalized by parents’ educational attainment and occupational status (von Stumm, Deary, & Hagger-Johnson, 2013), SES captures a multitude of factors that interact to shape children’s neurocognitive development through synergistic biological pathways (Jensen, Berens, & Nelson, 2017). In meta-analytic reviews, parents’ SES has been shown to account for about 9% of the variance in children’s educational achievement (Sirin, 2005; Strenze, 2007). In early childhood, children’s own intelligence test scores begin to predict educational achievement, accounting for about 5% of the variance (Honzik, Macfarlane, & Allen, 1948). By the school years, children’s previous educational achievement becomes the best predictor of their later educational and occupational outcomes. Educational achievement is highly stable throughout the school years, with year-to-year correlations of about .70, which means that earlier achievement accounts for about 50% of the variance of later achievement (Rimfeld et al., 2018).

In recent years, inherited DNA differences have been identified that can also predict educational achievement. For decades it has been known from twin and adoption studies that genetic differences taken as a whole account for most of the variance in educational achievement, with heritability estimates typically around 60% for all subjects at all ages (de Zeeuw, de Geus, & Boomsma, 2015; Rimfeld et al., 2018). Two breakthroughs in DNA research have made it possible to identify inherited DNA differences responsible for this high heritability (Plomin, 2018). The first began over a decade ago with the construction of ‘SNP chips’, arrays that are able to genotype inexpensively and quickly the most common type of inherited DNA difference, a difference in a single nucleotide base, called a single-nucleotide polymorphism (SNP).

The second breakthrough was to realise that the tiny effects of individual SNP associations from the GWA results for a trait, even those that do not individually meet the criterion of genome-wide significance, can be aggregated to create powerful predictors of a trait, called genome-wide polygenic scores (GPS) (Plomin & von Stumm, 2018). Because educational attainment (years of education) is obtained as a demographic marker in most GWA studies, it has been possible to include more than a million adults in a meta-analytic GWA study of educational attainment (Lee et al., 2018). A GPS derived from this GWA study predicts up to 13% of the variance in educational attainment in adults in independent samples (Lee et al., 2018). We have shown that a GPS for educational attainment predicts even more variance (15%) in tested educational achievement in 16-year-old adolescents (Allegrini et al., 2018), making this the most predictive GPS for any behavioural trait at present.

Although the expression of genome is modifiable and influenced by environmental factors, inherited DNA differences do not change throughout life. It follows that a GPS can predict 16-year-old educational achievement just as well at birth as it can at age 16. The ability to predict from birth is shared with family SES but not with other predictors such as children’s intelligence or educational achievement. GPS are completely reliable in the sense of test-retest reliability, and they are free from the measurement errors that typically affect data from psychometric tests and self-reports.

Although GPS and SES may be thought of, respectively, as quintessential indices of nature and nurture, their prediction estimates in fact capture both genetic and environmental effects (Kong et al., 2018; Krapohl & Plomin, 2016; Selzam et al., 2019). Prediction from GPS includes passive gene-environment correlations (prGE) that emerge because parents create a family environment that corresponds to their genotypes and, by extension, also correlates with the genotypes of their offspring. Because prGE stem from the parents’ genotype, they are genetic in origin but environmentally mediated (Selzam et al., 2019). Similarly, SES is often assumed to represent solely environmental advantages of wealth and privilege (Conley & Fletcher, 2017), but it is actually just as heritable as most other complex traits, with estimates from twin studies of about 50% (Branigan, McCallum, & Freese, 2013; Polderman et al., 2015). The main ingredients in most SES scores are parents’ educational attainment and occupational status, both of which are substantially heritable (Tambs, Sundet, Magnus, & Berg, 1989; Taubman, 1976). Indeed, adult educational attainment, specifically the number of years spent in formal education, is the target trait for the GWA analysis used to create the GPS that predicts children’s educational achievement. Therefore, it is not surprising that the prediction of educational achievement from family SES is in part mediated by GPS (Belsky et al., 2016; Belsky et al., 2018; Krapohl & Plomin, 2016; Selzam et al., 2019).

Previous research has suggested that the prediction of educational achievement from both GPS and SES increases during development (Allegrini et al., 2018; Selzam et al., 2017; von Stumm, 2017), but their relative contribution over time to educational achievement or their interaction have not been explicitly tested before. Longitudinal latent growth curve analysis, which disentangles initial effects on a developmental measure (intercept) from systematic increases and decreases which follow (slope), is particularly useful for this purpose (McArdle, 2009). A growth curve model also allows testing for interaction effects of GPS and SES, thereby evaluating the gene-by-environment interaction hypothesis that the influence of genetic factors on cognitive development are weaker in lower-SES families (Gottschling et al., 2019; Tucker-Drob & Bates, 2016).

## METHODS

### Sample

The present sample is drawn from the Twins Early Development Study (TEDS), a longitudinal twin study that recruited more than 10,000 twin pairs born in England and Wales in 1994 through 1996. The recruitment process and the sample are described in detail elsewhere (Rimfeld et al., 2019). The TEDS sample was at its inception representative of the UK population in comparison with census data and remains considerably representative, despite some attrition (Rimfeld et al., 2019). After excluding twins who experienced severe medical complications during the first two years of life, the analyses sample included 4,890 individuals of European ancestry from TEDS for whom genotype data and information on socioeconomic status and educational achievement from age 7 through to 16 were available. This selection includes children from TEDS, who are typically developing and comparable in gestational stage, birth weight, and maltreatment experience. The analysis sample is slightly more advantaged in terms of SES (mean = 0.14, SD = 0.95) than the full TEDS sample at conception (mean = 0, SD = 1). The unadjusted GPS for years spent in education differed minimally between the analysis sample (mean = 3.12, SD = .28) and the all TEDS participants with genotype data (mean = 3.11, SD = .28). Our analysis sample included one twin from each monozygotic twin pair and all dizygotic twin pairs. We also confirmed our analyses and findings in a smaller sample, which included only one twin from each twin pair (N =3,297) for maximal independence of data (see Table S2 for descriptive statistics in both samples, and Table S3 for modelling results in the smaller sample.)

### Measures

#### Genome-wide Polygenic Scores (GPS)

Saliva and buccal cheek swab samples were collected, DNA was extracted and genotyped to compute genome-wide polygenic scores (GPS) based on the latest genome-wide association (GWA) study for years of education, called educational attainment (EA3) (Lee et al., 2018). Overall, 7,363,646 genotyped and well-imputed SNPs were retained for the analyses after stringent quality control, but to ease high computational demands of the software LDpred (Vilhjálmsson et al., 2015) for polygenic scoring in large samples, we further excluded SNPs with info <1, leaving 515,100 SNPs for analysis (see Supplementary Methods for details.) GPS were constructed using the Bayesian LDpred approach, which estimates causal effect sizes from GWA summary statistics by assuming a prior for the genetic architecture and LD information from a reference panel (Vilhjálmsson et al., 2015). Here, GPS are constructed as the weighted sum of trait-increasing alleles for each unrelated genotyped individual in the TEDS sample using a causal fraction of 1, which assumes that all SNPs contribute to the development of the trait. The GPS were adjusted for the first ten principal components of the genotype data, chip, batch and plate effects using the regression method and the standardized residuals were used for all subsequent analyses (see Supplementary Methods S2 for details).

#### Educational achievement

In the UK, compulsory schooling follows a national curriculum that is organised into blocks of years called ‘key stages’ (KS). At the end of each key stage, teachers formally assessed each child’s performance. The twins’ teachers recorded their grades in English and mathematics in the National Pupil Database (NPD) following the UK National Curriculum guidelines, which are formulated by the National Foundation for Educational Research (NFER; http://www.nfer.ac.uk/index.cfm) and the Qualifications and Curriculum Authority (QCA; http://www.qca.org.uk). KS1 through KS4 were recorded in the NPD at age 7, 11,14 and 16, respectively. The English grades assessed students’ reading, writing, and speaking and listening; the mathematics grades referred to knowledge of numbers, shapes, space, and to using and applying mathematics and measures. At age 16, twins’ grades in English and mathematics were awarded for subtests of the General Certificate of Secondary Education (GCSE) exam, a standardized examination taken at the end of compulsory schooling in the UK, and recorded in the NPD. For each key stage, we built composite scores of children’s NPD grades for English and mathematics.

#### Socioeconomic status (SES)

Families’ SES was assessed at the first contact, when the twins were 1.5 years old, with parents reporting their educational qualifications, their occupational positions and the twins’ mother’s age on the birth of first child. A composite measure of SES (Krapohl & Plomin, 2015) was calculated taking the mean of the standardized scores for mothers’ and fathers’ educational level, mothers’ and fathers’ occupational status, and mothers’ age at birth of the first child.

### Statistical analysis

Students’ educational achievement at each key stage was adjusted for age and gender, and standardized regression residuals were used in all subsequent analyses. After testing correlations across all study variables, we fitted a series of latent growth curve (LGC) models using the R package Lavaan (Rosseel, 2012). LGC models differentiate two components of individual differences in growth. The first component, known as the intercept, reflects individual differences that are evident at the first assessment of a trait, in our case, performance in KS1 at age 7, and remain stable across assessment waves (i.e., from KS1 through KS4). The second component, known as slope, captures individual differences in the rate of change that occurs over time in the same trait, here change in educational achievement from KS1 through KS4.

In line with LGC modelling conventions, loadings for the intercept factor were set at 1, while factor loadings for the slope represented the number of KS (i.e. slope loadings were set at 0,1, 2, and 3), thereby defining the starting point of the slope at KS1. We first tested two- and three-factor LGC models (i.e. intercept, slope and quadratic factor, which was specified by the squared slope loadings, i.e. 0,1, 4, and 9) in order to identify which best represented the data. We then tested the predictive validity of EA3 and SES for educational achievement at KS1 (i.e. intercept factor) and for change in school performance from KS1 through KS4 (i.e. slope factor). We also tested if EA3 and SES interacted in the prediction of educational achievement over time.

## RESULTS

The prediction of educational achievement from both GPS and SES increased steadily throughout the school years, as shown in Figure 1. From ages 7 to 16 years, the association of GPS with educational achievement increased from .22 to .36 (i.e. from 5% to 14% of the variance) and that of SES from .31 to .48 (i.e. from 10% to 23% of the variance). The substantial gains in the predictive power of GPS and SES from age 7 to age 16 is particularly noteworthy because educational achievement was highly stable across the school years, with longitudinal correlations ranging from .61 to .87 (Supplementary Table S1). GPS and SES were correlated at .35, which reflects genetic influences on SES, in the sense that SES is partially heritable, as well as influences of SES on GPS, in the sense that SES partly mediates the effects of GPS on educational achievement, as discussed earlier.

**Figure 1.**
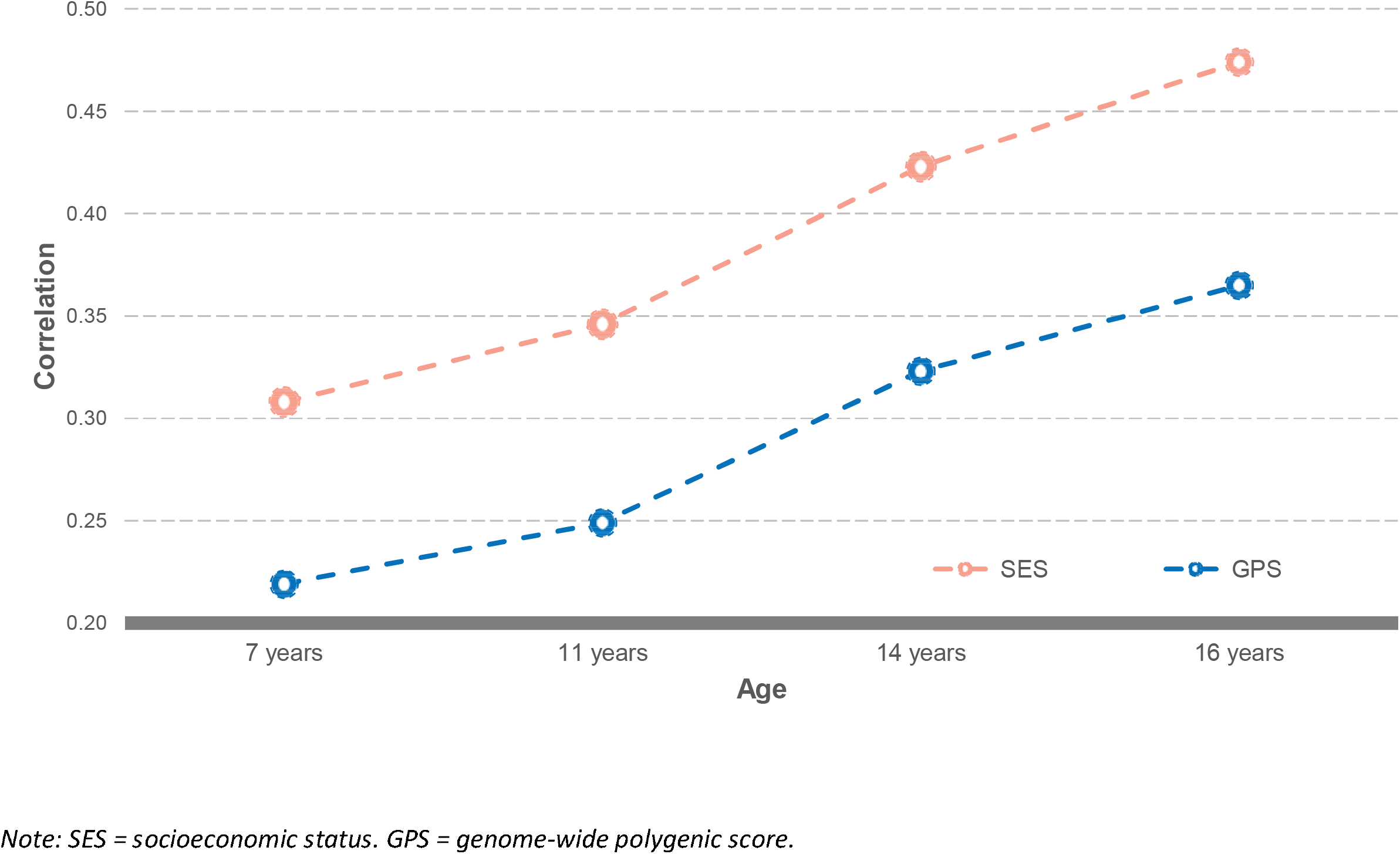
GPS and SES correlations with educational achievement at 7, 11,14 and 16 years

The potential practical implication of the magnitude of these predictions is especially apparent at the extremes. Dividing the sample into 10 equal-sized groups (deciles) from low to high GPS and similarly from low to high SES, Figure 2 depicts the mean standardised educational achievement (GCSE) scores at age 16. The lowest and highest deciles differed by more than 1.2 SD for GPS and 1.5 SD for SES. These average GCSE scores for the lowest and highest deciles translate into grades of C+ and A-, respectively, for GPS, and C- and A for SES. It is also noteworthy that the association of both GPS and SES with educational achievement was linear.

**Figure 2.**
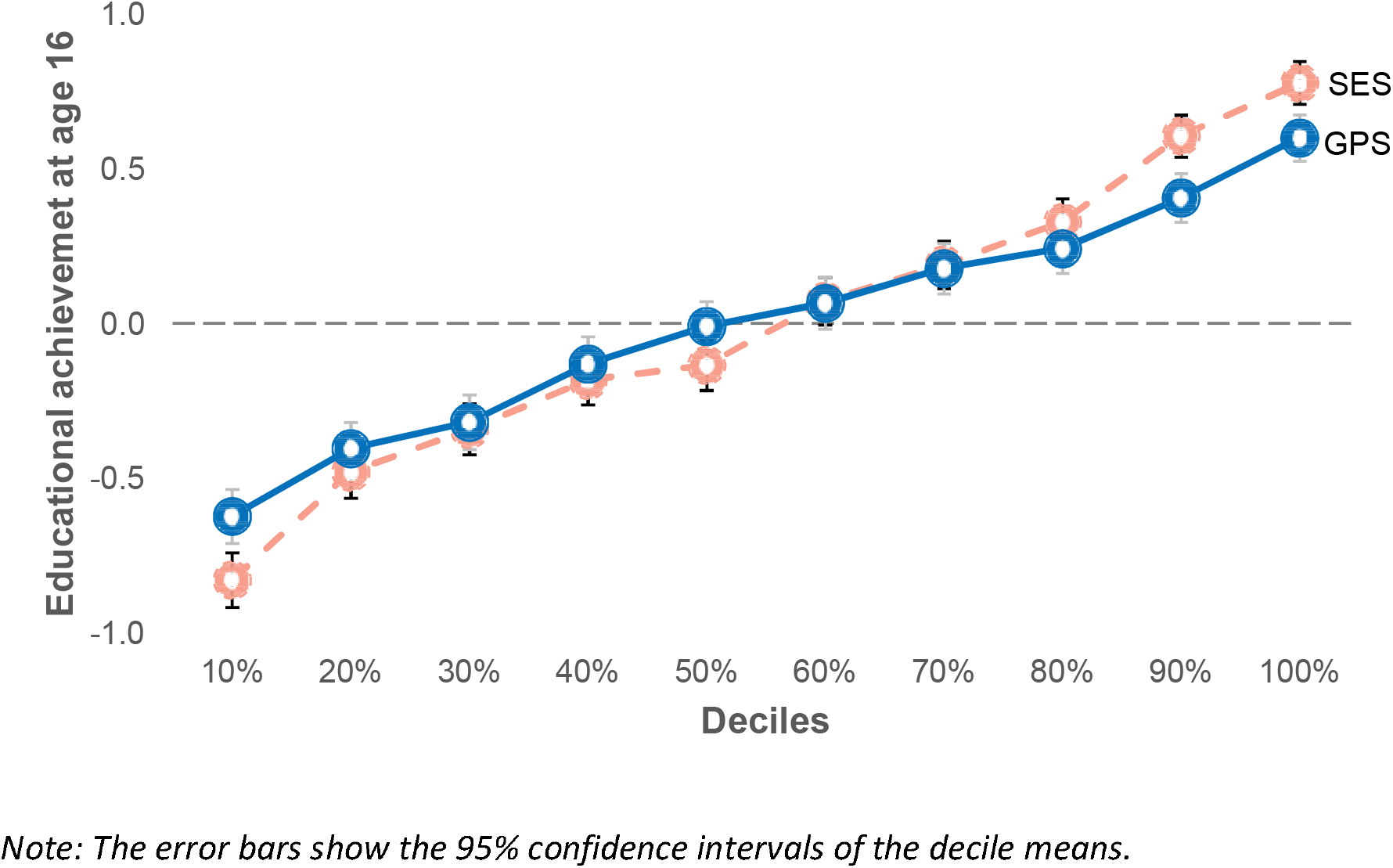
GPS and SES decile means for standardised educational achievement scores at age 16

The comparatively large mean differences across the decile levels of the predictors mask a considerable range of individual differences. Figure 3 shows the substantial overlap between the distributions of scores within the lowest and highest deciles for GPS and for SES (i.e. grey area). For example, 9% of the individuals in the lowest GPS decile had higher educational achievement scores than the mean score of the highest GPS decile (Figure 3a). Conversely, 10% of those in the highest GPS decile had lower educational achievement scores than the mean of the lowest GPS decile. For SES, the analogous overlap is 5% and 3%, respectively (Figure 3b).

The 95% confidence intervals for the decile means in Figure 2 indicate that accurate estimates of educational achievement can be made on average for extreme groups. However, the overlap in distributions in Figure 3 highlights the limits of prediction at the level of individuals.

**Figure 3.**
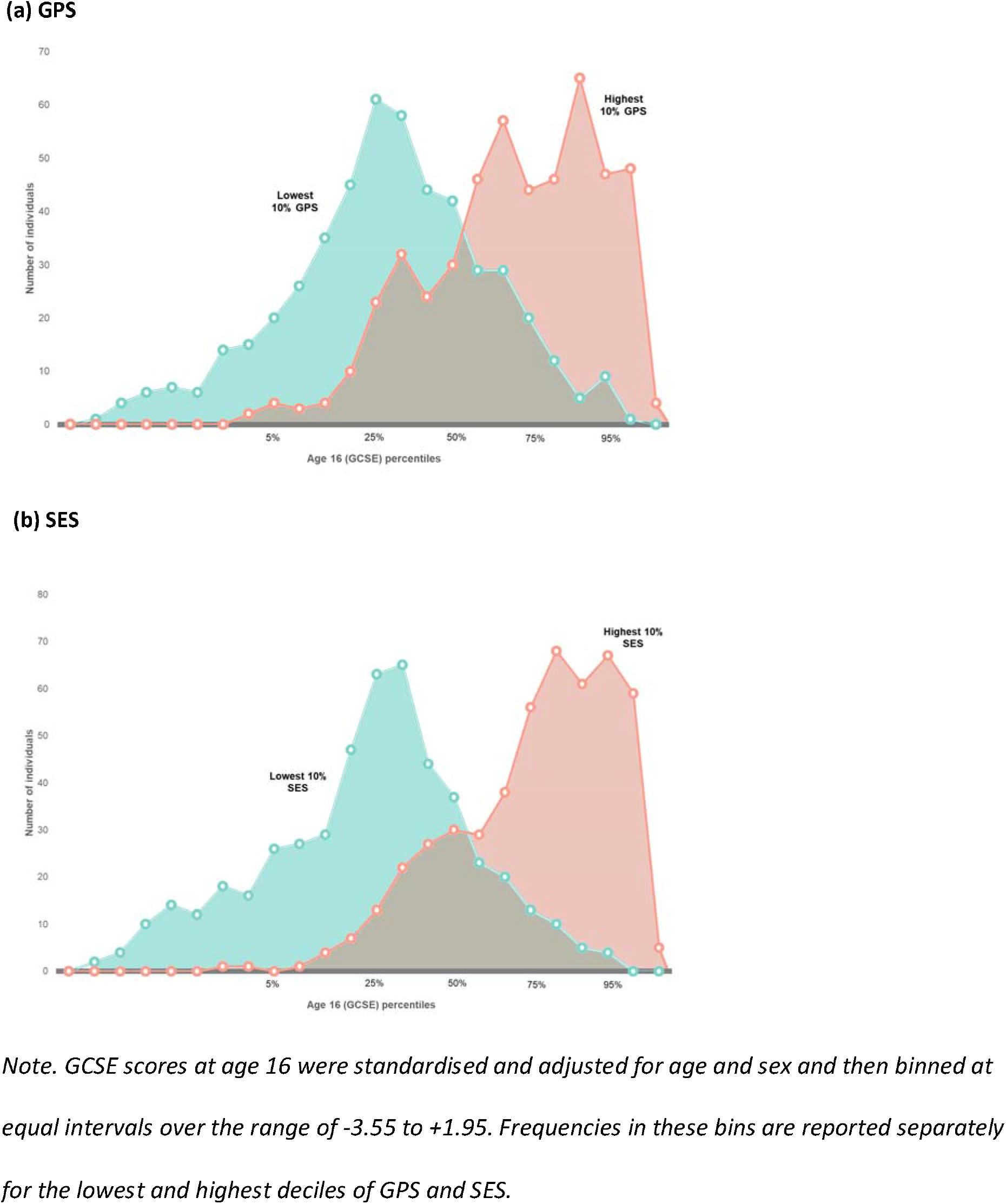
Overlap in distributions of educational achievement (GCSE) at age 16 for lowest and highest deciles of (a) GPS and (b) SES

Our LGC analysis indicated that a two-factor model including an intercept and a linear growth factor (i.e. slope) fit worse than a three-factor model with an additional quadratic factor (χ^2^_diff_ (4) = 150.23, p < .001). However, because the quadratic factor correlated -.86 with the slope factor, we retained the more parsimonious two-factor solution that differentiated latent factors representing individual differences in intercept and in slope for educational achievement (model fit: CFI = 0.990; RMSEA = 0.077, 90% confidence interval = 0. 067 to 0.088). Intercept and slope factors had, respectively, variances of .676 (SE = .022) and .038 (SE = .002), and they correlated r = .08 (p = .018). Their means were 0, a consequence of adjusting the educational achievement measures for age and gender (i.e. standardized residuals).

We used GPS, SES and their interaction as joint predictors of the two LGC factors of educational achievement from KS1 through KS4. GPS and SES were significantly and independently associated with latent growth factors representing individual differences in intercept and slope of educational achievement, as reported in Table 1. In other words, both GPS and SES independently contributed to the prediction of individual differences educational achievement that were observable at the beginning of school and remained stable over time (intercept) but also to systematic increases and decreases (slope) in achievement across the school years.

**Table 1:**
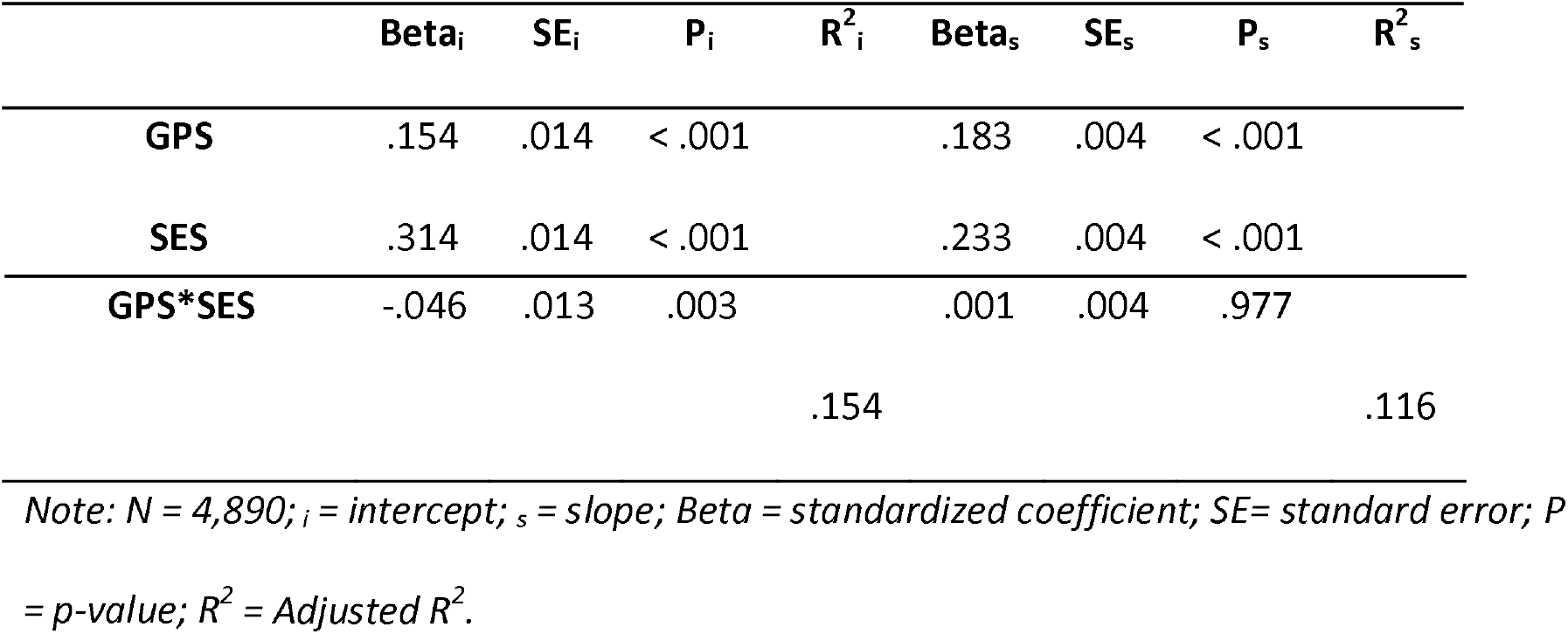
Parameter estimates for GPS, SES and their interaction as predictors of latent growth factors (intercept and slope) of educational achievement

Table 1 also shows that the effects of GPS and SES interacted in the predictions of the intercept but not for the slope. These results might seem to support the gene-by-environment interaction hypothesis that genetic influences on cognitive development vary across different levels of family SES (Gottschling et al., 2019; Tucker-Drob & Bates, 2016). However, the negative beta of the interaction term for the intercept is in fact a result in the opposite direction than originally proposed (Tucker-Drob & Bates, 2016): It appears here that GPS scores were more predictive of differences in educational achievement at age 7 in children from low SES families than in those from high SES backgrounds. That said, the effect size associated with the interaction effect was small (i.e. 0.2% of the variance; see also our additional analyses in Supplementary Materials), suggesting that the influence of GPS and SES on educational achievement is primarily additive.

Together, GPS and SES accounted for 15.4% of the variance of the intercept and 11.6% of the slope, as indicated by R^2^ in Table 1. The independent effects of GPS and SES, which we estimated by summing the squared betas in Table 1, contributed 12.3% (i.e., 2.4% + 9.9%) to the variance of the intercept and 8.7% (i.e., 3.3% + 5.4%) to the variance of the slope. The difference between R^2^ and these independent contributions of GPS and SES represents overlapping prediction by GPS and SES: 3.2% for the intercept and 2.8% for slope.

## DISCUSSION

The two best predictors of children’s educational achievement that are available from birth are parents’ SES and children’s inherited DNA differences (i.e. GPS). The current analyses showed that SES and GPS account together for 27% of children’s differences in educational achievement across the course of compulsory schooling. Furthermore, SES was a stronger predictor of educational achievement from age 7 through to 16, accounting independently for 15.3% of the variance. By comparison, GPS accounted independently for 5.7% of the variance. Also, 6% of the variance in education achievement from age 7 through to 16 could be attributed to the shared influence of SES and GPS (i.e. common variance).

As noted earlier, SES and GPS capture both genetic and environmental effects (Krapohl & Plomin, 2016; Kong et al., 2018; Selzam et al., 2019). Our findings shed light on predictive associations between SES, GPS and educational achievement but they do not allow causal inferences about genetic and environmental influences. Even so, the accurate prediction of educational achievement is an essential component of identifying those children that suffer the greatest risk for poor educational outcomes. These children are likely to have the greatest need for and potential benefit from intervention programmes that seek to improve educational achievement. Furthermore, we speculate that this group may be instrumental for future research that elucidates the gene-to-behaviour pathways and the mechanisms that underlie the associations between SES, GPS, and educational achievement. We caution, however, that our findings may be specific to samples of European ancestry because extensive GWA studies, which are required for identifying DNA variants that are reliably associated with a target phenotype, are at this time not available in populations with other ancestries (Mills & Rahal, 2019). Furthermore, we acknowledge that the assessment of SES in our study is coarse and only captures some of its many elements (Jensen et al., 2017).

Analyses of the extremes of GPS and SES highlight our developmental findings. We identified four extreme groups in our sample using cut-offs of ± 1SD in SES and GPS (Figure 4; see also Supplementary Figure S1). Children with high GPS from high SES family backgrounds and those with low GPS from low SES families (i.e. high-high and low-low groups, respectively) differed significantly in achievement at age 7 and the achievement gap steadily widened between the groups throughout the school years. By age 16, the mean educational achievement of the two groups differed by 1.9 SD, which is equivalent to a grade of A-in the high-high group and C-in the low-low group. In the high-high group, 77% went on to attend university at age 18 as compared to 21% in the low-low group.

**Figure 4.**
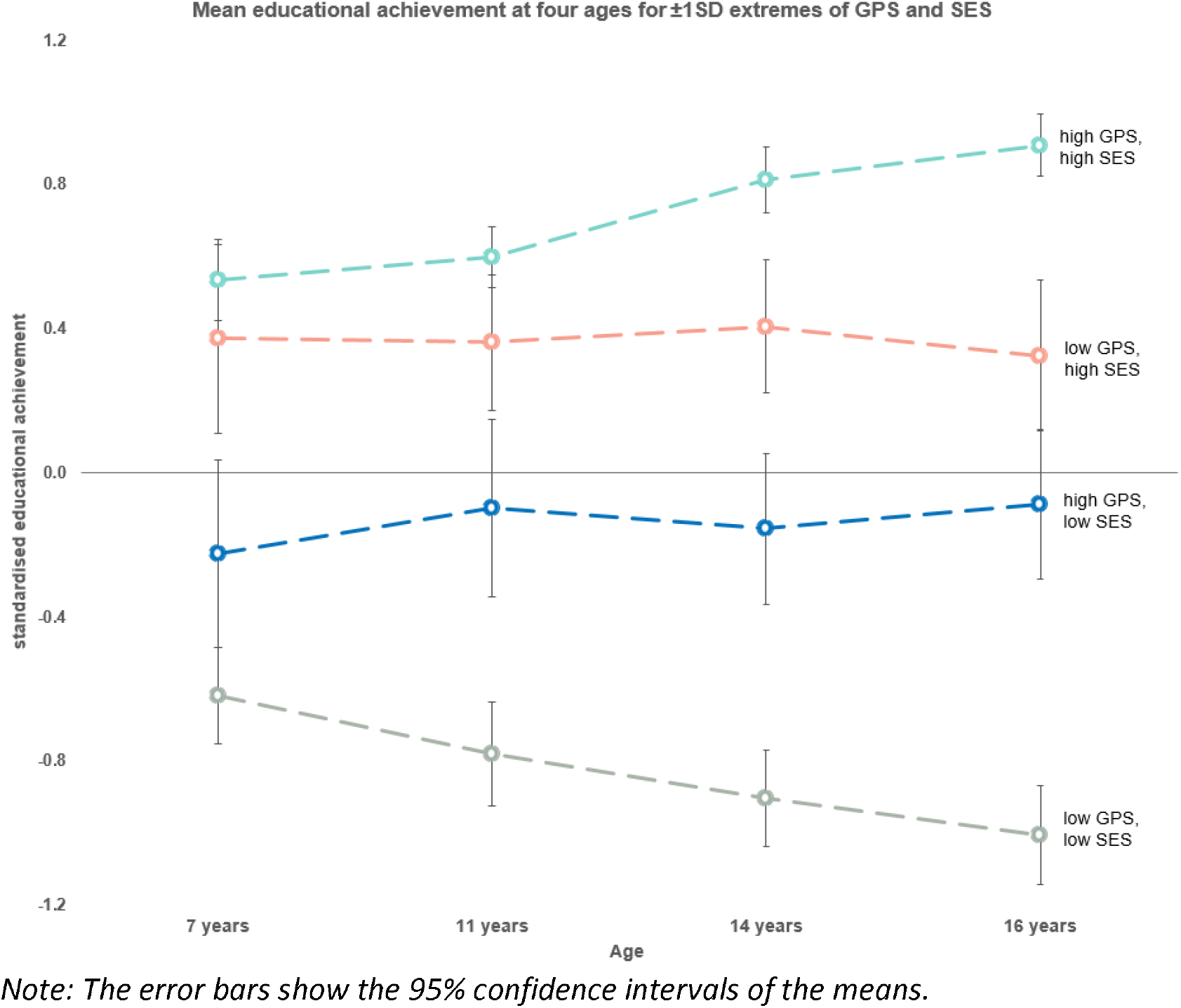
Mean educational achievement at four ages for ± 1SD extremes of GPS and SES

We found that GPS contributed even more to children’s differences in growth in educational achievement (3.3% of the variance) as it did to their differences at the beginning of compulsory schooling (2.4%). This is likely to explain why heritability is not much reduced by attempts to produce fairer measures of ‘added value’ of schools (more recently called measures of ‘progress’ in the UK), which statistically adjust current educational achievement for cognitive ability and earlier achievement (Haworth, Asbury, Dale, & Plomin, 2011). Specifically, we speculate that correcting for ability and earlier achievement does not in fact correct for the genetic contribution to growth, which is independent of intercepts (von Stumm, 2017).

By comparison, our analyses showed that SES contributed less to the growth in educational achievement (5.4%) than it did to children’s performance differences that are stable throughout compulsory schooling (9.9%). This finding suggests that measures of ‘added value’ of schools do a better job of controlling for the effects of SES than they do for GPS. This interpretation is supported by twin analyses that found that estimates of shared environmental influence, which include the effects of SES, are halved using measures of added value (Haworth et al., 2011).

Turning to the two middle, ‘mixed’ groups in Figure 4, it appears that high GPS may help compensate for disadvantages in school performance in children from low SES families. Our analyses showed that at age 16, children with high GPS from low SES backgrounds had mean GCSE scores that were close to average (-.09 standard score) and 47% of these children went on to attend university, as compared to only 21% in the low-low group. We note, however, that to an even greater extent high SES helped compensate for the disadvantage of low GPS children. Their mean GCSE scores were slightly above average (.32 standard score) and they were much more likely to later go to university (62%). Nonetheless, these predictive effects of GPS and SES were largely independent and additive (see Supplementary Figure S2). That is, we found no evidence for an interaction between genes (GPS) and environment (SES), in line with a meta-analysis on this issue (Tucker-Drob & Bates, 2016).

## CONCLUSIONS

Our major finding is that SES and inherited DNA differences aggregated in GPS are powerful predictors of educational achievement, accounting together for 27% of children’s differences in achievement across the course of compulsory schooling. The influence of GPS and SES is particularly dramatic at the extremes of the distribution. We suggested, for example, that high GPS partially compensates for the disadvantages of children from low-SES families, increasing their chances of going to university from 21% to 47%. This raises the possibility of doing more to help this group reach its full potential. Nonetheless, the substantial overlap between the distributions of scores within the lowest and highest deciles for GPS and SES indicates the limits of prediction at the level of individual students.

The potential application of predictive capacity of the kind demonstrated here will require complex decision-making. The basis for those decisions goes beyond purely scientific criteria to issues of ethics and social values. Papers like the present one provide an essential empirical grounding for discussion. It is our hope that our results and others like them can serve to open doors for individual children, not close them, by stimulating the development and provision of personalized environments that can appropriately enhance, supplement, and remediate educational achievement.

