## Supplementary materials for "Predicting educational achievement from genomic measures and socioeconomic status"

**Title**

**Table of Contents**

Table S1. Correlations between GPS, SES and educational achievement from age 7 through 16 years

Figure S1. Scatterplot (r = .30) between standardized GPS and SES showing ± 1SD cut-offs for extremes analyses.

Figure S2. The effects of GPS and SES on GCSE scores (KS4) show no evidence for interaction between genes (GPS) and environment (SES).

Methods S1. Genotyping and quality control

Methods S2. Creating polygenic scores using LDpred

Methods S3. Latent growth curve model results including only one twin from each dizygotic twin pair

Table S2. Descriptive statistics for full sample and for sample including only one twin per pair

Table S3. Latent growth curve parameter estimates for GPS, SES and their interaction as predictors of latent growth factors (intercept and slope) of educational achievement in the sample including one twin per pair

Table S1: Correlations between GPS, SES and educational achievement from age 7 through 16 years

|  | GPS | SES | Age 7 | Age 11 | Age 14 |
| --- | --- | --- | --- | --- | --- |
| GPS | - |  |  |  |  |
| SES | .35 | - |  |  |  |
| Age 7 | .20 | .31 | - |  |  |
| Age 11 | .24 | .35 | .66 | - |  |
| Age 14 | .32 | .42 | .63 | .78 | - |
| Age 16 | .36 | .48 | .61 | .74 | .87 |

*Note: N = 4,890 for all variables*

Figure S1. Scatterplot (r = .35) between standardized GPS and SES showing ± 1SD cut-offs for extremes analyses.

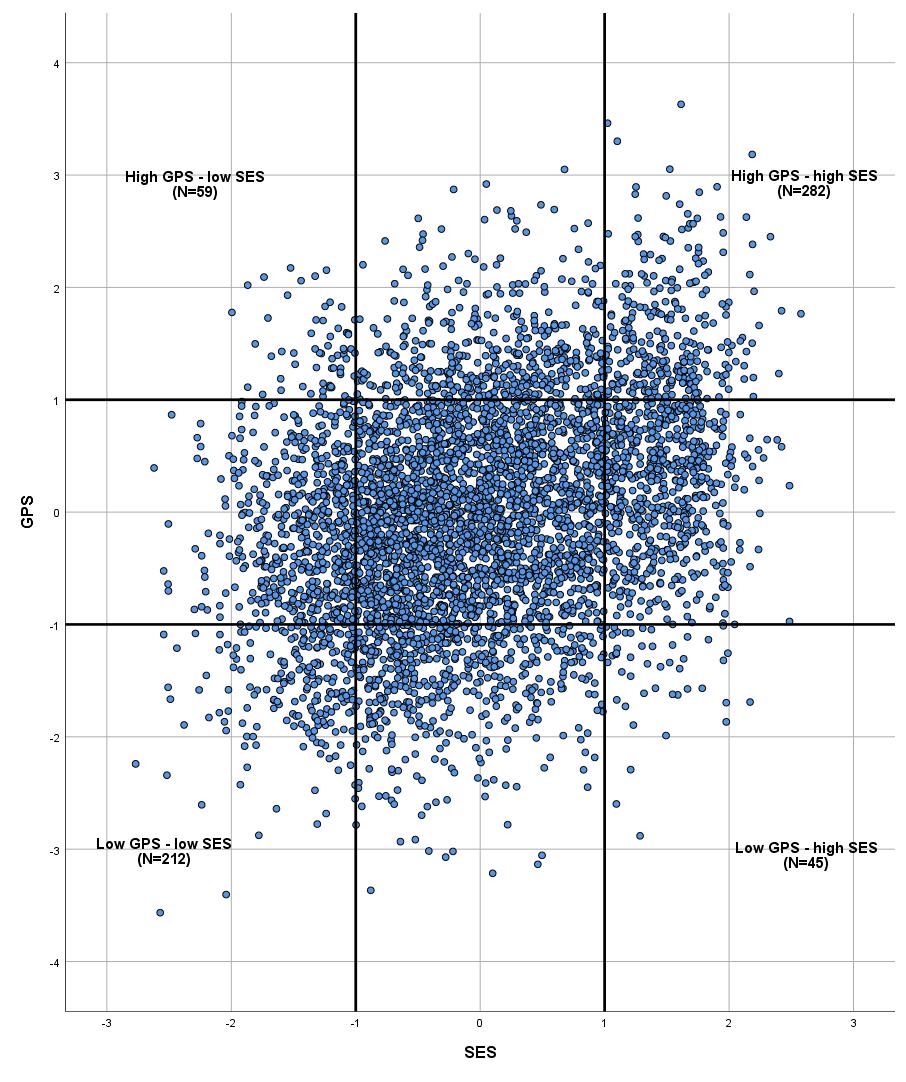

*Note: N = 4890. GPS and SES are standardized scores.*

*The bold lines indicate ± 1SD cut-offs.*

Figure S2. The effects of GPS and SES on GCSE scores (KS4) show no evidence for interaction between genes (GPS) and environment (SES).

GCSE Standardised Scores

|  | Low GPS | High GPS |
| --- | --- | --- |
| Low SES | -1.00486 | -0.0879284 |
| High SES | 0.324626 | 0.908877 |

**Methods S1. Genotyping and quality control**

DNA from 8,122 individuals was extracted from saliva and buccal cheek swab samples and hybridized to HumanOmniExpressExome-8v1.2 genotyping arrays at the Institute of Psychiatry, Psychology and Neuroscience Genomics & Biomarker Core Facility. Raw image data were pre-processed in GenomeStudio according to Illumina Exome Chip SOP v1.4. (<http://confluence.brc.iop.kcl.ac.uk:8090/display/PUB/Production+Version%3A+Illumina+Exome+Chip+SOP+v1.4>). Prior to genotype calling, 919 multimapping SNPs and 501 samples with call rate <0.95 were removed. Following initial QC, the program ZCALL was used to augment genotype calling.

DNA from 3,747 individuals was extracted from buccal cheek swabs and genotyped at Affymetrix, Santa Clara, California, USA. From the extracted DNA samples, 3,665 samples were successfully hybridized to AffymetrixGeneChip 6.0 SNP genotyping arrays ([http://www.affymetrix.com/support/technical/datasheets/genomewide_snp6_datashe et.pdf](http://www.affymetrix.com/support/technical/datasheets/genomewide_snp6_datashe%20et.pdf)) using experimental protocols recommended by the manufacturer.

Raw image data were pre-processed at the Wellcome Trust Sanger Institute, Hinxton, UK for genotyping as part of the Wellcome Trust Case Control Consortium 2 (<https://www.wtccc.org.uk/ccc2/>). All pre-processing was conducted according to the manufacturer’s guidelines (http://www.affymetrix.com/support/downloads/manuals/genomewidesnp6_manual.pdf). Following initial QC, the program CHIAMO was used for genotype calling (https://mathgen.stats.ox.ac.uk/genetics_software/chiamo/chiamo.html).

After initial quality control, the same quality control was performed on samples from each of the platforms (Illumina and Affymetrix) separately using PLINK (Purcell et al., 2007), R(Team, 2015) and BCFtools(Li, 2011) and EIGENSOFT(Patterson, Price, & Reich, 2006; Price et al., 2006).

DNA samples were excluded from subsequent analyses on the basis of call rate (<0.98), suspected non-European ancestry, the presence of severe medical or psychiatry problems or severe medical complications during early gestation and relatedness other than dizygotic twin status. SNPs were excluded if the minor allele frequency was <0.5%, if more than 2% of genotype data were missing, or if the Hardy Weinberg *p*-value was lower than 10^-5^. Non-autosomal markers and indels were also removed. Associations between SNPs and the platform, batch, plate or well on which samples were genotyped were calculated; SNPs with an effect p-value < 10-4 were excluded.

A total sample of 10,346 samples, including 7,026 unrelated individuals; 3,320 additional individuals had a genotyped dizygotic co-twin. Genotype data following quality control were available for 4,776 individuals and 559,772 SNPs from the illumine array and 2,250 individuals and 635,269 SNPs from the Affymetrix array.

Genomewide genotypes from the two arrays were separately phased using EAGLE2 (Loh et al., 2016) and imputed using the Haplotype Reference Consortium (McCarthy et al., 2016) using the Positional Burrows-Wheeler Transform method (Durbin, 2014) and the imputation software Minimac3 1.0.13 (Fuchsberger, Abecasis, & Hinds, 2015), which are available from the Michigan Imputation Server (https://imputationserver .sph.umich.edu). A series of quality checks were performed before merging data from the two arrays, and variants with info <0.75 were excluded and SNPs that were non-overlapping between platforms were removed.

After merging, minor allele frequency differences were tested for between platforms and SNPS with an effect p-value <10^-4^ were removed. Those SNPs with a Hardy-Weinberg p-value >10^-5^ were also removed. Following these criteria, 7,363,646 genotyped and well-imputed SNPs were retained for analyses. Only unrelated individuals were included in the present analyses. To ease high computational demands of the software LDpred (Vilhjálmsson et al., 2015) for polygenic scoring in large samples, we further excluded SNPs with info <1, leaving 515,100 SNPs for analysis.

**Methods S2. Creating polygenic scores using LDpred**

Genomewide polygenic scores were calculated using the Bayesian approach, LDpred, which has been shown to outperform predictive accuracy of the conventional clumping and p-value thresholding approach (Vilhjálmsson et al., 2015). Here, a posterior effect size is derived for each SNP by re-weighting the original summary statistic coefficient by the relative influence of a SNP given its level of linkage disequilibrium with surrounding SNPS and a prior on the effect size of each SNP. The prior is based on the heritability of the trait and the fraction of markers assumed to casually influence the trait. GPS is then calculated as the sum of the trait-increasing alleles weighted by their posterior effect size estimate. Unlike the conventional clumping and thresholding approach, LDpred retains all SNPs common between GWA summary statistics and genotype data in the target sample.

For the present study we applied a causal fraction of 1, which assumes that all SNPs contribute to the development of the trait. Due to the high computational demand of LDpred, especially in larger sample sizes including many SNPs, we made further restrictions to our analyses, only including the 515,100 SNPs that were perfectly imputed (info score of 1) to reduce analytical load. Only genotypes of unrelated individuals were used to estimate LD structure in our sample because levels of LD are considerably higher in relatives compared to unrelated individuals (Vattikuti, Guo, & Chow, 2012).

**Methods S3: Latent growth curve model results including only one twin from each dizygotic twin pair**

We replicated our latent growth curve analysis selecting only one twin from each dizygotic twin pair, resulting in an overall sample of 3,297 participants, compared to 4890 participants in the analyses reported in the main manuscript.

Table S2 shows the descriptive statistics for the study variables in both samples. We normed GPS in both samples by transforming scores to standardized z-scores (mean = 0; SD = 1).

Table S2: Descriptive statistics for full sample and for sample including only one twin per pair

|  |  | N | Mean | SD | Skew | Kurtosis |
| --- | --- | --- | --- | --- | --- | --- |
| Full analysis sample (main manuscript) | | | |  |  |  |
|  | SES | 4890 | 0 | 1 | 0.11 | -0.72 |
|  | GPS | 4890 | 3.12 | 0.28 | 0.01 | -0.03 |
|  | KS1 | 4890 | 0 | 1 | -0.09 | 0.03 |
|  | KS2 | 4890 | 0 | 1 | -0.88 | 0.78 |
|  | KS3 | 4890 | 0 | 1 | -0.48 | 0.03 |
|  | KS4 | 4890 | 0 | 1 | -0.43 | 0.02 |
| Sample including one twin per pair (SOM) | | | |  |  |  |
|  | SES | 3297 | -0.03 | 1 | 0.12 | -0.71 |
|  | GPS | 3297 | 3.12 | 0.28 | 0.03 | 0.00 |
|  | KS1 | 3297 | -0.03 | 1.01 | -0.11 | 0.08 |
|  | KS2 | 3297 | -0.02 | 1.01 | -0.86 | 0.67 |
|  | KS3 | 3297 | -0.02 | 1.01 | -0.48 | 0.03 |
|  | KS4 | 3297 | -0.01 | 1 | -0.43 | -0.01 |

A two-factor model including slope and intercept fit better than a one-factor model (χ^2^_diff_ (3) = 433.05, p < .001) but worse than a three-factor model (χ^2^_diff_ (4) = 105. 11, p < .001). However, the third factor, a curvilinear factor, correlated -.88 with the slope factor. For this reason, we retained the two-factor solution (model fit: CFI = 0.990; RMSEA = 0.078, 90% confidence interval = 0.066 to 0.091).

Associations of GPS and SES with latent growth factors are reported in Table S3. The parameter estimates differ marginally from those achieved in the sample including both DZ twins that are reported in the main manuscript. However, the p-values are affected by the sample size change, with the interaction between GPS and SES in the prediction of the intercept being associated with a p-value of .022 in the smaller sample but with a p-value of .003 in the bigger sample. The effect sizes associated with the interaction effect were 0.02% and 0.01% of the variance in educational achievement in the bigger and smaller sample, respectively. By contrast, the other parameter's significance and effect size remained largely unchanged across bot samples. We interpret these findings to show that the identified interaction effect is at best weak, if not spurious altogether, and that the effects of SES and GPS on educational achievement are primarily additive.

Table S3: Latent growth curve parameter estimates for GPS, SES and their interaction as predictors of latent growth factors (intercept and slope) of educational achievement in the sample including one twin per pair

|  | *Beta_i_* | *SE_i_* | *P_i_* | *R^2^_i_* | *Beta_s_* | *SE_s_* | *P_s_* | *R^2^_s_* |
| --- | --- | --- | --- | --- | --- | --- | --- | --- |
| *GPS* | *.135* | *.017* | *< .001* |  | *.178* | *.005* | *< .001* |  |
| *SES* | *.317* | *.017* | *< .001* |  | *.232* | *.005* | *< .001* |  |
| *GPS*SES* | *-.035* | *.015* | *.022* |  | *.011* | *.005* | *.633* |  |
|  |  |  |  | *.148* |  |  |  | *.115* |

*Note: _i_ = intercept; _s_ = slope; Beta = standardized coefficient; SE= standard error; P= p-value*
